# PERFECT PCR: Advancing DNA Data Storage to Near-Maximal Density

**DOI:** 10.64898/2025.12.15.694532

**Authors:** Rushant Sabnis, Han Zhang, Bingzhe Li, Arum Han, Paul de Figueiredo, Qing Sun

## Abstract

DNA-based data storage offers unprecedented storage density and durability compared to traditional media, but it faces challenges in long access latency and limited encoding efficiency. Current DNA storage methods achieve only 8% of the theoretical maximum storage density (TMSD) due to biological constraints and error rates. Here, we demonstrate an approach that improves both synthesis speed and encoding density, bringing DNA data storage closer to practical implementation. We describe Polymerase-Enabled Rapid and Frugal Error Correcting Technology (PERFECT) PCR, which integrates PCR with hyperthermophilic EndoNucS enzymes for efficient DNA mismatch detection and cleavage, enabling accurate and fast DNA synthesis using overlapping oligonucleotides. This approach reduced latency while streamlining the overall DNA synthesis process, achieving significantly lower error rates in amplicons compared to state-of-the-art methods, thereby reducing physical redundancy. Specifically, we achieved a substantially reduced error rate of 0.06 errors per kilobase, with 95% of sequences verified as accurate. As a result, our technique reached 94% TMSD, representing a twelve-fold improvement over current systems and approaching the theoretical limit for existing sequencing technologies. We further validated the versatility of PERFECT PCR by successfully encoding and retrieving digital images, music files, and multilingual text files with near-perfect information recovery. By addressing the core challenges of latency, density, and accuracy, our approach brings DNA-based data storage significantly closer to practical implementation. In addition, the accurate DNA synthesis methods described here may be applied to *de novo* genome assembly and therefore have broad implications for the synthesis and analysis of biological systems.

**SUMMARY:** Low cost, rapid, and error free *de novo* DNA synthesis for DNA-based data storage and gene synthesis applications.

## INTRODUCTION

DNA-based data storage has emerged as a promising alternative for storing and archiving large volumes of data (*1–22*). With global demand for data storage growing exponentially, the International Data Corporation (IDC) projects that the global data storage demand will reach 175 Zettabytes by 2025, approximately 6 times more than the demand in 2018 (*23, 24*). By 2040, an estimated 2.4 billion kilograms of high-grade silicon will be required to manufacture computer chips and storage device components, yet current trends suggest that only 1% of this demand can be met (*18*). DNA-based data storage systems have demonstrated the potential to achieve an impressive storage capacity of up to 214 petabytes per gram (Pbytes/g) of DNA, exceeding current storage technologies such as solid-state drives by approximately six orders of magnitude (*22*). In addition, DNA-based storage offers exceptional durability and is environmentally friendly (*25*). It requires significantly lower energy to maintain and does not rely on precious metals used in conventional storage devices (*26, 27*). DNA has a half-life 1,000 times longer than silicon-based devices, making it ideal for long-term storage and data retention (*6, 12, 23*). These characteristics position DNA as a promising solution for addressing the growing global demand for sustainable and scalable data archiving, particularly in light of increasing demands for data storage (*11, 28–30*).

Despite these attractive features, DNA-based data storage technology remains in its nascent stages and faces numerous challenges, particularly long latency, high error rates, and prohibitive monetary costs associated with *de novo* DNA synthesis (*22, 31*). These challenges have prevented DNA from being widely adopted in data storage systems. Although significant strides in computational methods have mitigated the need for physical redundancy and enhanced accuracy, predicting lost data accurately remains a challenge, particularly in cases where models are not sufficiently trained (*6, 22, 24, 25, 32, 33*). To address these challenges, synthesizing *de novo* DNA using overlapping oligonucleotides has proven to be faster and more cost-effective compared to synthesizing entire gene blocks from scratch (*34, 35*). This synthesis strategy involves assembling short, 40-150 base pair overlapping oligonucleotides, produced via phosphoramidite or enzymatic chemistry, into longer DNA fragments (*36–41*). However, the accuracy of the final products, defined as the ratio of the total number of molecules without a single error to the total number of molecules, depends heavily on the coupling efficiencies during oligonucleotide synthesis. The maximum practical phosphoramidite coupling efficiency reported to date is approximately 99.5%, which poses a significant limitation for modern gene synthesis applications, including DNA data storage. The coupling efficiency results in an error rate of 0.5% which translates into 5 errors per 1,000 base pairs synthesized, leading to 40% of the assembled sequences of size 1 kb containing at least one error. Consequently, only 60% of the sequences are error-free, representing the overall template accuracy. This level of accuracy is inadequate for the precision required in DNA-based data storage applications (*42–44*).

Currently, two main strategies have been employed to achieve high-accuracy DNA molecules. The first involves the removal of incorrectly synthesized sequences from the pool of correct ones, using methods such as High-Performance Liquid Chromatography (HPLC) and Polyacrylamide Gel Electrophoresis (PAGE) (*35*). These techniques were designed to selectively eliminate sequences containing significant errors, such as insertions and deletions (indel errors), thereby enriching the population of error-free molecules. However, such methods are laborious and time consuming. Alternatively, the second strategy focuses on enhancing the accuracy of the synthesis process itself, to minimize the generation of incorrect sequences.

Enzymatic error correction strategies, for example, target and repair mismatches within DNA (*45–51*). Researchers have used these enzymes to bind to and remove error-containing DNA, or cleave mismatched bases, followed by amplification of the corrected templates via PCR, enhancing the accuracy of synthesized DNA. While these approaches help reduce errors, complete elimination of inaccuracies through a single round of enzymatic error correction has shown low efficacy. Consequently, multi-step error correction combined with PCR has been explored and shown to further reduce the error rates of synthesized DNA (*34, 35*). However, the labor-intensive and time-consuming nature of multiple sequential PCR and error correction steps, coupled with the challenges of reusing the same set of primers, often leads to inefficient binding to PCR products. This outcome can result in nonspecific amplification or amplification failure, complicating further PCR rounds and limiting progress beyond a few cycles.

Our strategy integrates enzymatic error correction directly into the PCR amplification process. A major limitation of commercially available error correction enzymes was their inability to remain active under high temperatures and the frequent thermal cycling of PCR. To overcome this challenge, we identified error detection endonucleases from hyperthermophilic organisms, which remain stable and active at 98°C, the peak temperature during PCR. We then designed the Polymerase-Enabled Rapid and Frugal Error Correcting Technology (PERFECT) PCR approach, which represents an innovative integration of enzymatic DNA mismatch detection and cleavage within PCR thermal cycling (**Fig. 1**). Using this approach, we achieved low-error amplicons with an error rate of 0.06 errors/kb, which enabled the realization of 94% of the theoretical maximum storage density (TMSD), with a data retrieval success rate of 96.5%, representing a twelve-fold increase in storage density compared to current state-of-the-art DNA-based storage approaches. PERFECT PCR addresses the challenge of data loss by reducing the errors in sequences storing the data thereby enhancing the overall reliability of the stored data and supporting computational methods that reduce redundancy and improve accuracy.

**Fig. 1.**
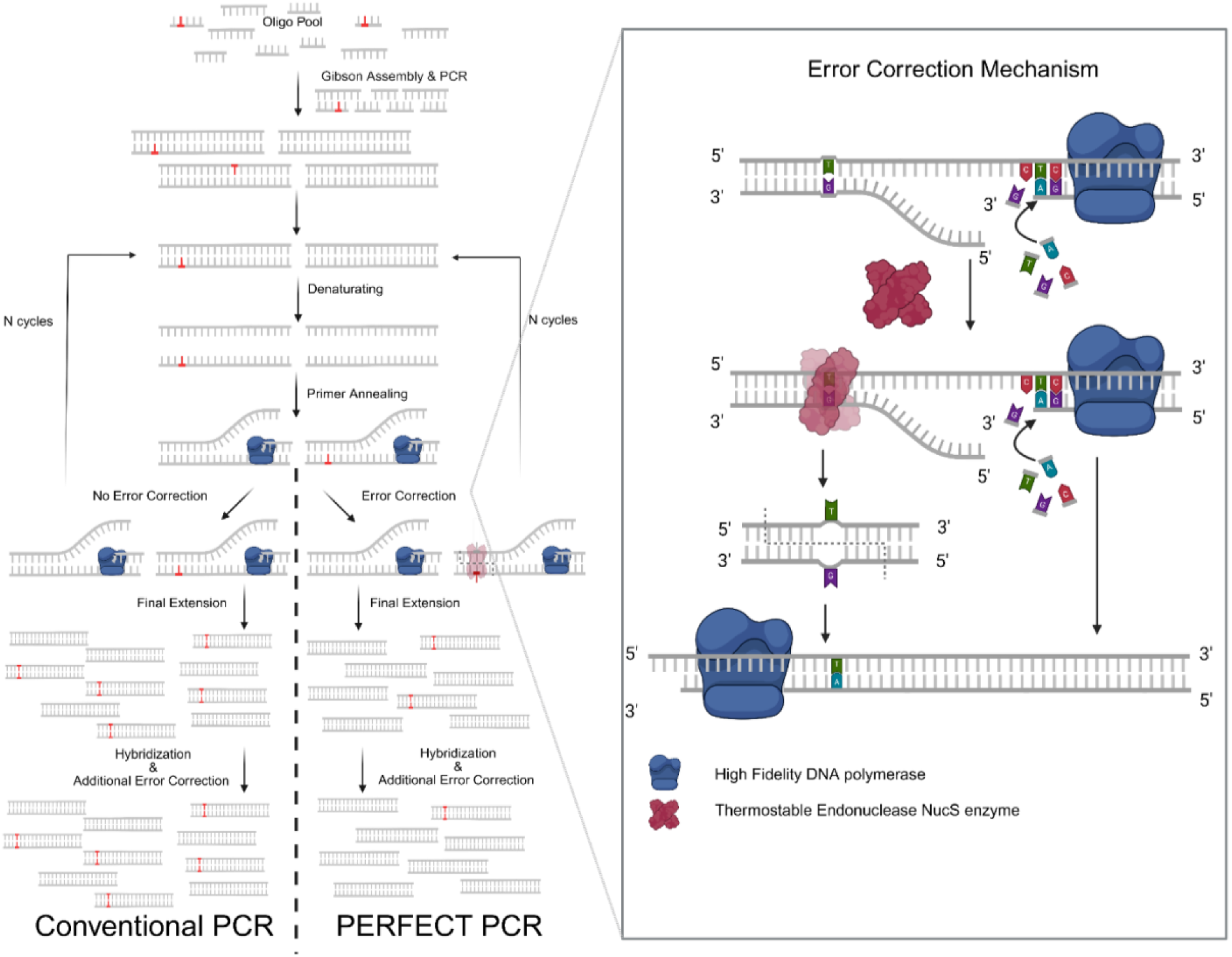
Schematic of PERFECT PCR and mechanism of error correction. In conventional PCR, errors introduced during amplification are propagated across successive cycles, leading to an accumulation of errors. In contrast, PERFECT PCR incorporates an error-correction enzyme that mitigates error propagation at each cycle. Additionally, the inclusion of a hybridization and reannealing step further reduces error rates resulting in a low error rate product.

## RESULTS

We aimed to develop a streamlined pipeline for the efficient synthesis of *de novo* DNA with a reduced error rate, facilitating the use of overlapping oligonucleotides for DNA-based data storage. Incorporating commercial enzymes, such as CorrectASE or surveyor nuclease (*34, 35, 52, 53*), into PCR components of the pipeline design was not feasible due to their inability to withstand the high temperatures present in PCR reactions. Moreover, these enzymes exhibited insufficient thermostability to withstand the repeated temperature fluctuations associated with PCR cycling, including the high temperature denaturing, primer annealing and DNA extension steps, resulting in rapid loss of activity and limiting their suitability for PCR-based applications. To address these challenges, we hypothesized that leveraging the heat stability of enzymes derived from hyperthermophilic microbes could facilitate their seamless integration into PCR reactions, correcting errors in oligonucleotides synthesized using phosphoramidite chemistry, ultimately generating error-free amplicons. Central to this approach was our focus on the mismatch-specific endonuclease EndoNucS, derived from hyperthermophilic Archaea. Previous studies characterized the EndoNucS enzyme from *Pyrococcus abyssi* (Pab), including the elucidation of its active site and functional mechanisms (*54, 55*). Additionally, EndoNucS from *Thermococcus kodakarensis* (Tko) was shown to be a functional homolog of the archaeal MutS/MutL system, which cleaves double-stranded DNA (*56*). Although EndoNucS from *Thermococcus kodakarensis* (Tko) was initially considered for these studies, experimental testing (**Fig. S1A**) confirmed that it did not maintain stability at temperatures above 85°C, limiting its utility for higher-temperature applications, including PCR.

To expand our search for thermostable mismatch-specific endonucleases, we used Basic Local Alignment Sequence Tools (BLAST) to identify EndoNucS enzymes with sequence similarities to EndoNucS-Tko (*57–66*). This analysis allowed us to categorize EndoNucS enzymes into two groups: Thermococcoaceae (comprising *Thermococcus*, *Pyrococcus*, and *Paleococcus*), and Methanomada (including *Methanobacter*, *Methanococci*, and *Methanopyri*), (**Fig. S2A**). After narrowing our search to the *Thermococcus* and *Pyrococcus* groups, which are typically thermophilic, we selected representative EndoNucS enzymes from *Pyrococcus kukulkanii* (Pkl), *Pyrococcus furiosus* (Pfu), and *Thermococcus barophilus* (TbaMP), as these Archaea thrive at temperatures exceeding 100°C (**Fig. 2A**). We hypothesized that the EndoNucS enzymes from these organisms were likely to exhibit activity at the elevated temperatures required for PCR.

**Fig. 2.**
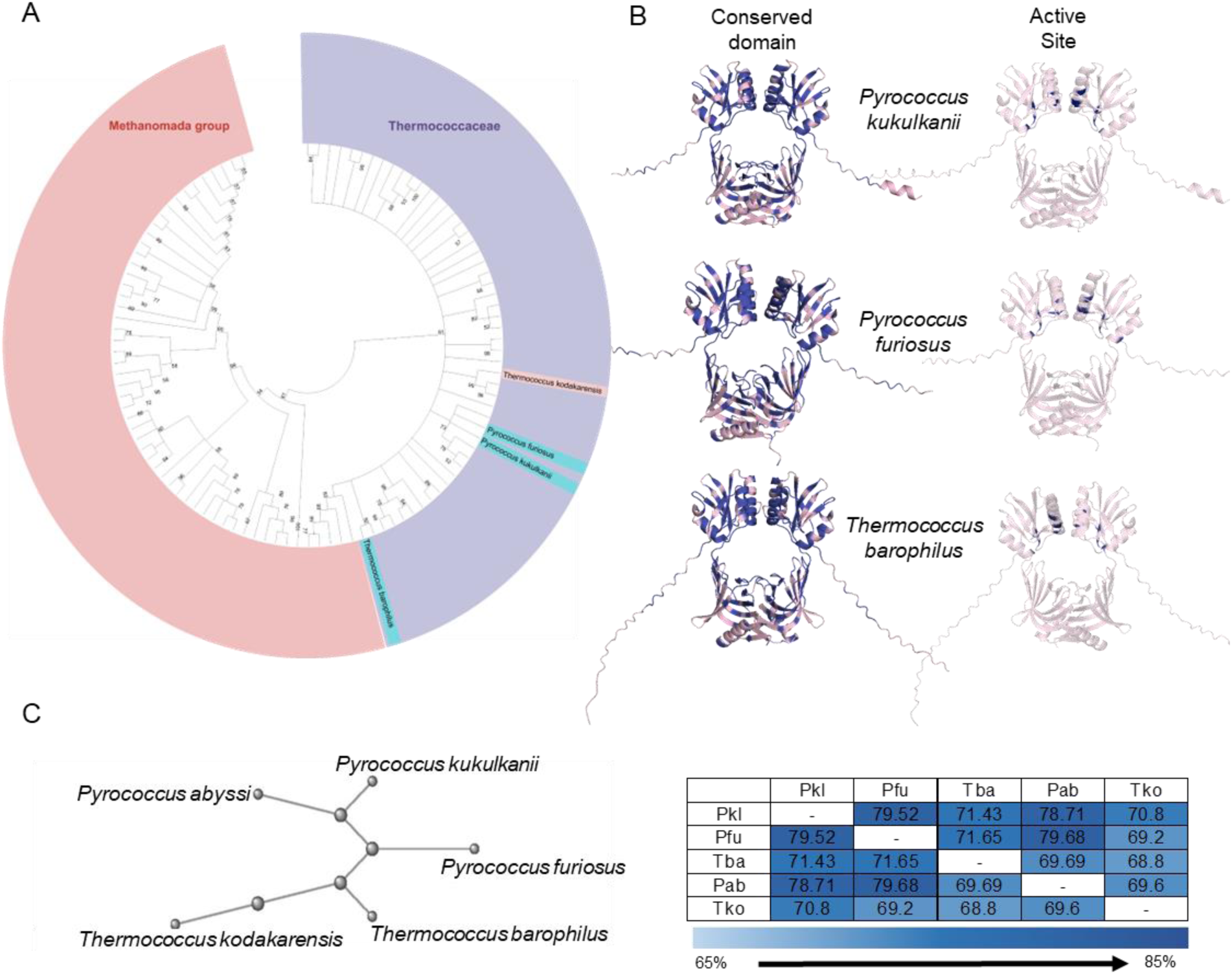
Phylogenetic Relationships, Structural Features, and Sequence Comparison of archaeal EndoNucS enzyme. (**A**) Phylogenetic tree of selected archaeal EndoNucS enzymes. *Thermococcus kodakarensis* (Tko), highlighted in red, represents the source organism of the enzyme characterized by Ishino et al. The species highlighted in cyan, *Pyrococcus kukulkanii* (Pkl), *Pyrococcus furiosus* (Pfu), and *Thermococcus barophilus* (Tba), denote the host organisms from which the EndoNucS homologs analyzed in this study were derived. Bootstrap values are indicated at each node to denote confidence in phylogenetic placement. (**B**) Predicted dimeric structures of EndoNucS enzymes from Pkl, Pfu, and Tba, generated using AlphaFold illustrating conserved domains and catalytic residues. Conserved regions and active site residues are indicated by dark, blue-colored amino acids. (**C**) Multiple sequence alignment of selected EndoNucS enzymes. The study focuses on enzymes from Pkl, Pfu, and Tba. For reference, *Pyrococcus abyssi* (Pab) represents the first characterized NucS enzyme, while *Thermococcus kodakarensis* (Tko) has been extensively studied in prior literature. All sequences share 68–80% pairwise identity.

We conducted an amino acid sequence comparison between EndoNucS-Pkl, EndoNucS-Pfu, and EndoNucS-TbaMP, alongside several enzymes described in the literature (*56, 67, 68*). Our analysis revealed significant conservation of the endonuclease active sites, with overall sequence similarity ranging from 65% to 85% (**Fig. 2C& Fig. S2C**). Additionally, 3D structural analysis using AlphaFold demonstrated that these enzymes exhibited strikingly similar folds (**Fig. S2B**). Structural modeling revealed that the interrogated enzymes formed dimers with a central hollow cavity, which served as the active site for cleaving mismatched DNA into two fragments. Enzyme-substrate interaction analysis, as shown in **Fig. 2B**, reveals that the enzyme contains a positively charged domain that interacts with the negatively charged mismatched DNA, facilitating accurate mismatch detection and cleavage (*69*). The essential catalytic sites and the conserved domain were predicted to be positioned around the hollow cavity where DNA mismatches are recognized and cleaved (**Fig. 2B**). Finally, because these enzymes are derived from hyperthermophilic archaea, which thrive at temperatures exceeding 100 °C and contain stabilizing polar residues, they are expected to maintain structural integrity and activity at high temperatures, including during PCR. Several features of the enzymes were characterized. First, we assessed their thermal stability and noted that EndoNucS-Pfu, EndoNucS-TbaMP, and EndoNucS-Pkl remained stable at temperatures as high as 98°C (**Fig. 3A to C** & **Fig. S1B**), which contrasted with EndoNucS-Tko, which denatured at 85°C (**Fig. S1A**). Second, we determined the substrate specificity of the enzymes (**Table S1**). For these experiments, we used the commercially available Surveyor nuclease enzyme as a control. Our results suggested that the examined EndoNucS enzymes detected and cleaved G/G, T/T, and G/A mismatched. However, they were unable to cleave C/C mismatches. These findings aligned with reports in the literature, where C/C mismatches proved challenging to correct (*70–72*). Finally, results from dose response (**Fig. 3D to I**) and reaction kinetic (**Fig. 3J to L** & **Fig. S3**) assays allowed us to define and optimize mismatch cleavage reactions.

**Fig. 3.**
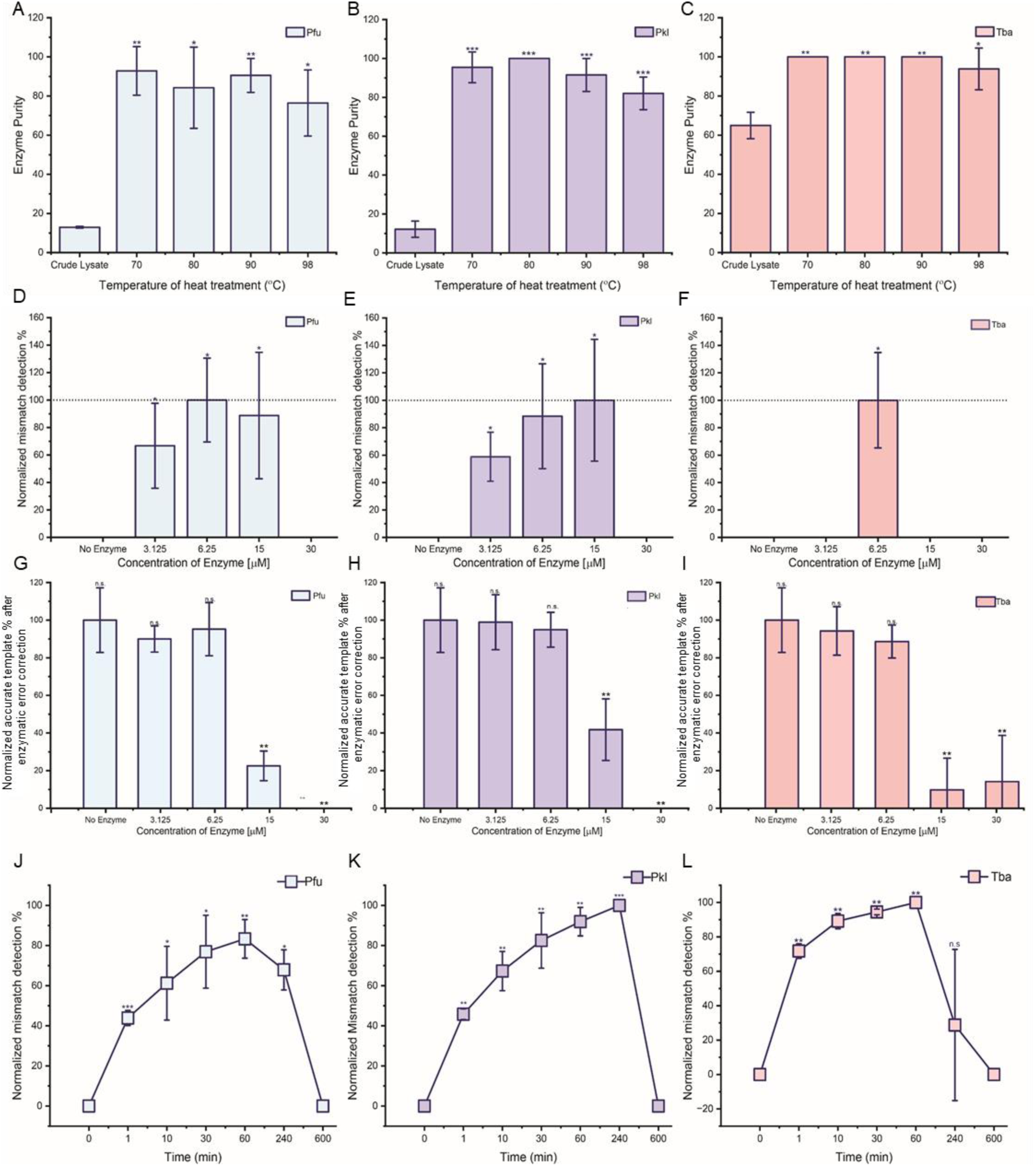
Hyperthermophilic archaea EndoNucS enzyme characterization. (**A-C**) Heat tolerance assay for Hyperthermophilic archaea EndoNucS Pfu, Pkl and Tba. (**D-F**) Concentration assay for Hyperthermophilic archaea EndoNucS Pfu, Pkl and Tba. (**G-I**) Exonuclease activity assay for hyperthermophilic archaea EndoNucS Pfu, Pkl and Tba. The figures demonstrate the extent of nonspecific and undesired exonuclease activity of the error correction enzyme. At higher concentrations, this activity becomes more prominent, leading to digestion of accurate templates. (**J-K**) Kinetic assay for Hyperthermophilic archaea EndoNucS Pfu, Pkl and Tba. n.s., p>0.05; *, p<0.05; **, p<0.01; ***, p<0.001.

We developed Polymerase-Enabled Rapid and Frugal Error Correcting Technology (PERFECT) PCR approach, which integrates enzymatic DNA mismatch detection and cleavage within PCR thermal cycling (**Fig. 1**). Acknowledging the limitations of single-round error correction, previous research explored up to two rounds of error correction, yielding significantly greater error reduction compared to a single round (*34, 35*). Unlike conventional PCR, which follows the standard steps of denaturation, primer annealing, and extension, the PERFECT PCR setup introduced error correction during the primer annealing step (**Fig. 1**). This methodological modification enabled error correction during each amplification cycle, resulting in at least 30 rounds of error correction throughout the PCR process (**Fig. 1**). We hypothesized that incorporating DNA mismatch cleavage directly during PCR cycling would enable multiple rounds of error correction, resulting in the generation of highly accurate amplified DNA.

For preliminary analysis, we synthesized the 717 bp green fluorescence protein (GFP) from overlapping oligo pools. We split the GFP protein sequence into 50 bp oligo pools with 25 bp overlap and then assembled these fragments using Polymerase Chain Assembly (PCA). The assembled product displayed 0.19 errors per kilobase (kb) (**Table S2**). After conducting PERFECT PCR using PCA products as template, PERFECT PCR with EndoNucS-Pkl exhibited the lowest error rate among the three tested enzymes, prompting our decision to employ it in subsequent experiments (**Table S2**). The EndoNucS-Pkl enzyme binds and cleaves mismatches on double-stranded DNA, thereby reducing the error rate of the amplified DNA product. In the PER-FECT PCR workflow, the thermostable enzyme functions during each round of thermal cycling to provide error correction. To further enhance accuracy, we modified the protocol by adding independent hybridization–error correction steps after completion of conventional PCR amplification. Due to its thermostability, EndoNucS-Pkl remains active at elevated temperatures and continues to recognize and cleave mismatches during these post-amplification reannealing steps. These extra hybridization-correction steps enhanced DNA rearrangement, promoting the reshuffling and formation of mismatches that could then be cleaved by EndoNucS. As a result, the error rate was reduced to 0.06 errors/kb, making it comparable to the lowest error rates reported in the literature (**Table 1& Table S3**) (*34*). Our data suggest that the improved performance primarily results from enzymatic mismatch elimination, as the additional hybridization-correction steps further enhance DNA accuracy. Further, errors in the PERFECT PCR product primarily stemmed from insertions and deletions of bases, while errors related to mismatches were promptly cleaved and corrected (**Table S2 and S4**). A comparison to prior reports indicated that, whereas most existing methods primarily correct indel-related errors, targeted correction of substitution-based errors during de novo DNA synthesis had not been previously reported (**Table S5**). To further evaluate our results, we directly compared our approach to a reported two-step error correction-PCR strategy using a commercially available CorrectASE enzyme (*35*). CorrectASE was selected as a benchmark due to its demonstrated capacity to achieve minimal residual error rates following two consecutive rounds of enzymatic correction (*35*), thereby serving as a stringent standard for evaluating error suppression efficiency. We found that PERFECT PCR had lower error rates, fewer steps, and was less time-consuming compared to the reported two-step error CorrectASE strategy (**Table 1, S6, Fig. 4& Fig. S5**).

**Fig. 4.**
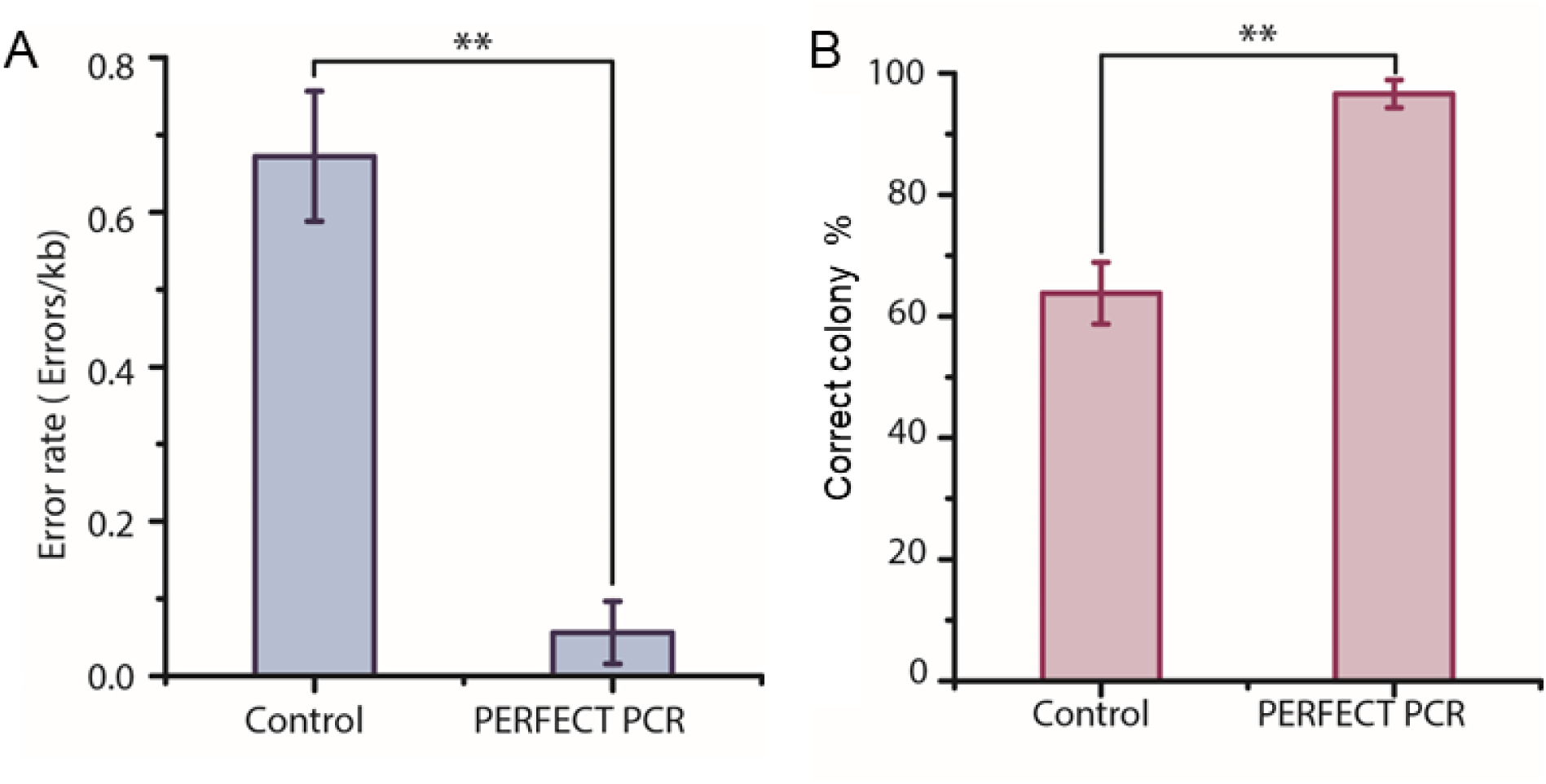
Comparison between conventional PCR and PERFECT PCR. (**A**) Error rate of GFP DNA sequence using conventional PCR (control) and PERFECT PCR after 35 PCR cycles. (**B**) Correct colony (%) using conventional PCR (control) and PERFECT PCR after 35 PCR cycles. (n=4; **p<0.01)

**Table 1.**
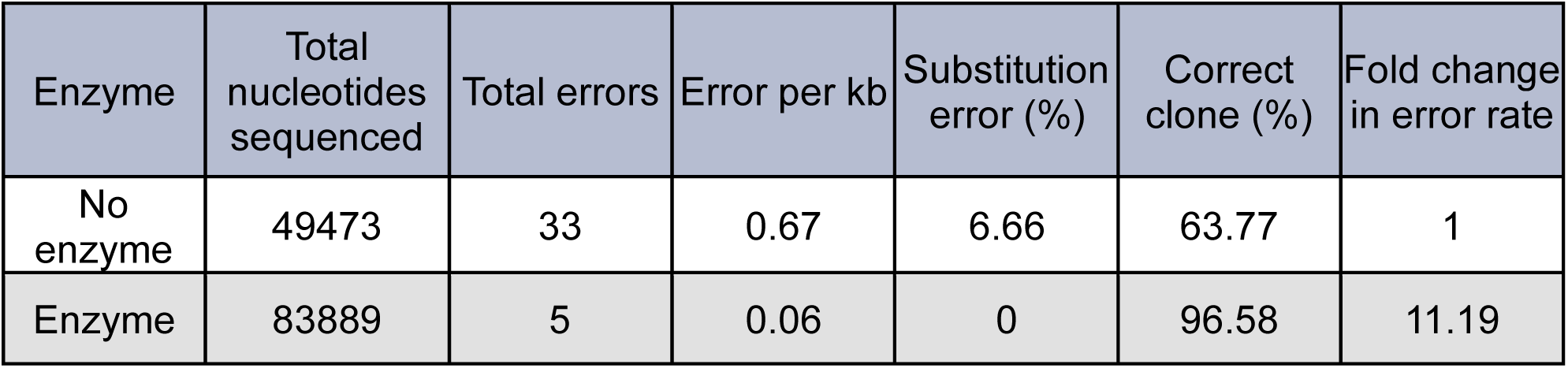
Comparison of conventional PCR and PERFECT PCR. The error rate reduced 11.19 folds from 0.67 to 0.06 errors/kb using the EndoNucS-Pkl enzyme. The number of accurate clones increases from 64% to 97%. This data was obtained by synthesizing eGFP gene using overlapping oligonucleotides. Correct clone % is defined as the ratio of the colonies without any errors to total number of colonies sequenced. The fold change in error rate was calculated by taking a ratio of the error rate without enzyme to error rate with enzyme. (Note: Compiled data from 4 biological replicates).

We tested the hypothesis that PERFECT PCR could be deployed for DNA-based data storage. First, utilizing the PERFECT PCR protocol, the accuracy of the GFP sequence DNA synthesized improved from 63% to 96%, measured as the proportion of error-free sequences relative to the total. This outcome yielded a storage density of 1.88 bits per nucleotide (**Fig. S6**), equating to 94% of the Shannon limit, the highest theoretically attainable storage density. Second, we compared this density to the state of art (0.16 bits/nucleotide) and found that the PERFECT PCR approach achieved a twelve-fold higher density (**Fig. 5A& Table S7**) (*33*). Third, we evaluated the ability of our approach to store image data. We implemented a delta-based encoding strategy with GC content optimization to convert a digital image into a DNA sequence. Using this approach, the “Eye of Tiger” image (88 × 96 pixels; 8,448 total pixels) was encoded into a 101,376-base-pair (bp) DNA sequence. For synthesis, the DNA was partitioned into 140 fragments of 720 bp and one terminal fragment of 576 bp and each fragment was flanked by universal forward and reverse primer binding sites to facilitate amplification. Oligonucleotide pools (oPools) were generated accordingly (**Fig. 5B**), with sequence data and oPools design parameters provided in the Supplementary Information. Experimental validation through DNA synthesis, high through-put sequencing, and subsequent error rate quantification demonstrated a marked reduction in substitution and indel frequencies with PERFECT PCR, lowering the error rate from 2.34 errors per kb to 0.31 errors per kb. Compared to our PER-FECT PCR system, enzymatic mismatch correction using Surveyor Nuclease and CorrectASE achieved error rates of 0.85 and 0.61 errors per kb, respectively (**Fig. 5C**). To assess the functional impact of these error profiles, we reconstructed images based on the corresponding error rates. Images derived from sequences treated with Surveyor Nuclease and CorrectASE exhibited significant visual degradation, whereas those processed with PERFECT PCR displayed superior visual clarity and preserved high image integrity (**Fig. 5D**). Lastly, we simulated the impact of error accumulation across multiple rounds of DNA-based data storage cycles. Each storage cycle was defined as one complete process of encoding data into DNA and subsequently retrieving it. Using the accuracy obtained from the GFP sequence experiment after PERFECT PCR, our simulations indicated that the non-error-corrected image exhibited a 92% error rate after just five cycles, leading to complete image degradation, whereas the error-corrected image accumulated fewer than 10% errors (**Fig. 5E& Fig. S4**).

**Fig. 5.**
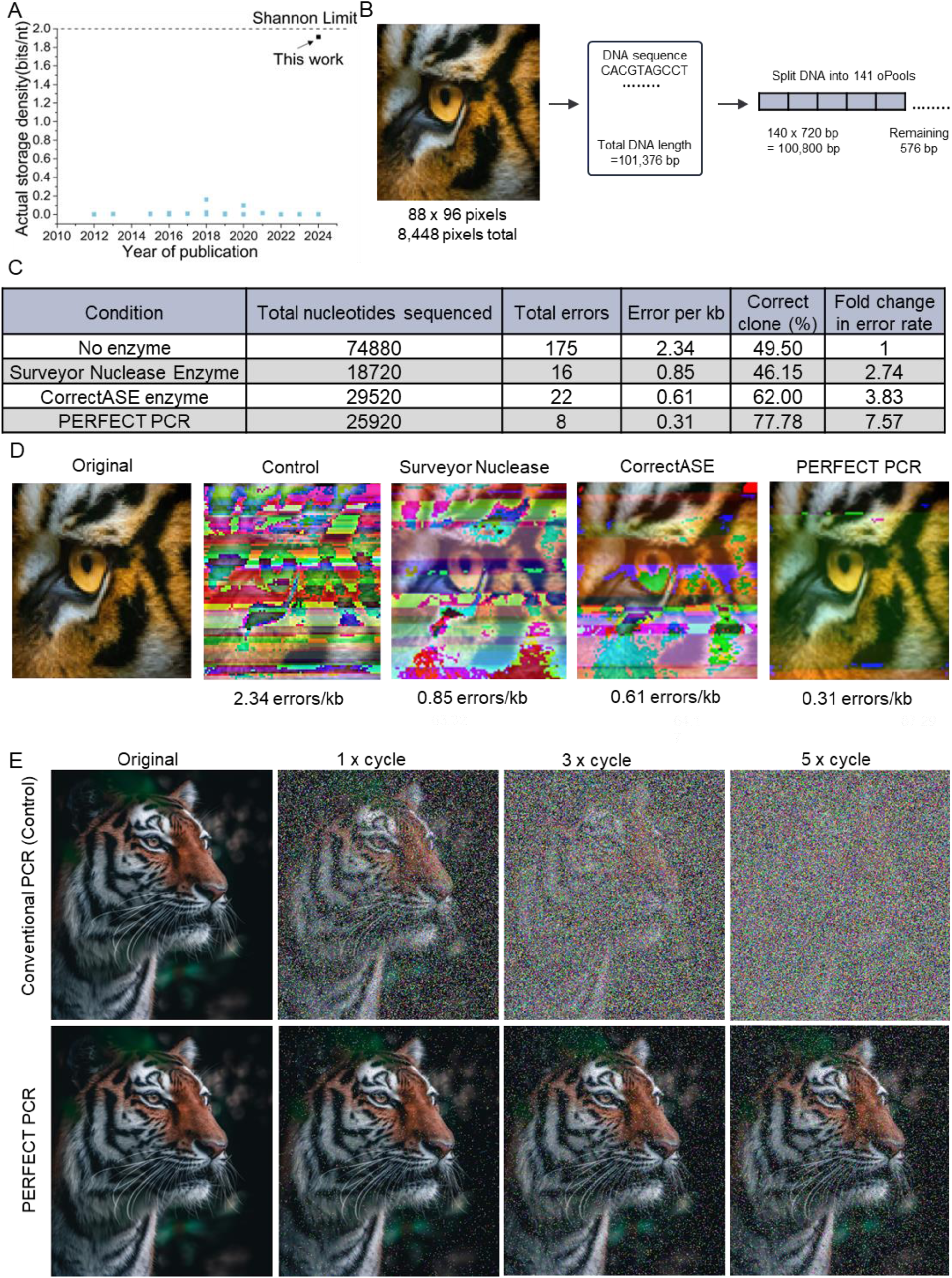
High-Density and Error-Free DNA-based data storage using PERFECT PCR. (**A**) Comparative analysis of storage densities achieved by PERFECT PCR and previously reported DNA data storage methods. After accounting for both physical and logical redundancy, PERFECT PCR achieves a storage density of 1.88 bits/kb, representing a 12-fold improvement over the next best state-of-the-art DNA data storage method. (**B**) Schematic representation of the DNA-based encoding and storage workflow for the Eye of the Tiger image. The input image (88 × 96 pixels; 8,448 pixels in total) was encoded into a 101,376 bp DNA sequence using a delta encoding algorithm in conjunction with GC-content balancing to enhance sequence stability. The resulting sequence was segmented into 141 overlapping oligonucleotides for synthesis, comprising 140 fragments of 720 bp and one terminal fragment of 576 bp. These oligonucleotides were synthesized as a pooled library (oPool) and subsequently amplified using the PERFECT PCR method to enable high-density and error-free data retrieval (**C**) Evaluation of error rates (expressed as errors per kb), correct colony % or retrieval accuracy (defined as the ratio of error-free colonies to total colonies), and fold-change in error rates with and without enzymatic error correction. Commercial enzymes tested include Surveyor Nuclease and CorrectASE. PERFECT PCR used the Pkl enzyme to represent the error correction strategy reported in this study. (**D**) Image reconstruction following DNA amplification using various PCR and error-correction strategies. The original image serves as a reference. The “Control” represents data recovered using conventional PCR. “Surveyor Nuclease” and “CorrectASE” refer to data recovered using the respective representative commercial error-correction enzymes. “PERFECT PCR” denotes data recovered using the method developed in this study. (**E**) Assessment of image recovery fidelity following multiple cycles of amplification. Images show progressive degradation after 1, 3, and 5 cycles of conventional PCR, with corresponding error rates of 37%, 60%, and 92%. In contrast, amplification using PERFECT PCR across the same number of cycles resulted in significantly lower error rates of 3%, 7.5%, and 9.9%, respectively. Simulated image reconstructions were generated based on experimentally determined error profiles from sequencing the eGFP control sequence.

We further demonstrated the versatility of PERFECT PCR by encoding and retrieving both musical compositions and multilingual text in DNA. Music files, including Spirit of Aggieland, Ode to Joy, and popular nursery rhymes, were encoded as note–duration pairs, while greetings in 14 languages from the Golden Record (*73*) were encoded character by character. After error correction, the music file was fully recovered with no corruption (**Supplementary Music File Data**) and the text file remained fully legible with some minor errors (**Table S13**). These results highlight PERFECT PCR as a robust strategy to store diverse data types in DNA with near-perfect information recovery.

## DISCUSSION

We developed PERFECT PCR, a novel method of reducing error rates in *de novo* DNA synthesized using overlapping oligonucleotides. We studied EndoNucS enzymes, including EndoNucS-Pfu, EndoNucS-Pkl, and EndoNucS-TbaMP, from Archaea that function by flipping and cleavage of mismatched DNA at the site of the error, generating a 5’ overhang. We observed that EndoNucS-Pfu, EndoNucS-Pkl, and EndoNucS-TbaMP remained stable at temperatures up to 98°C and maintained their mismatch-cleaving activity even at elevated temperatures of 55°C. We applied thermostable EndoNucS to develop the PERFECT PCR system for DNA synthesis by integrating them into PCR cycles. We achieved an error rate of 0.06 errors/kb and an accuracy of 96%, surpassing the state-of-the-art error correction method using CorrectASE/ErrASE enzyme (*34*), which has an error rate of 0.35 errors/kb and an accuracy of 80%. Next, we demonstrated the utility of PERFECT PCR for DNA-based data storage, where the approach significantly enhanced the quality of stored image data. We also used PERFECT PCR to store music and multilingual text files to demonstrate its versatility.

The low error rate associated with PERFECT PCR reduces the need for physical redundancy in data storage systems, minimizing the number of duplicate DNA sequences required to compensate for errors and thus enhancing storage density. While computational methods have advanced significantly, offering algorithms to mitigate biological limitations—particularly by improving accuracy through physical redundancy, they are still limited in accurately predicting lost data, especially when models are insufficiently trained. PERFECT PCR addresses this challenge by preventing data loss at the source, thereby improving the overall efficiency of DNA-based storage. When combined with advanced storage techniques like DNA Fountain and the use of DNA composite letters, logical redundancy—additional non-coded sequences added for error correction—can also be reduced, further optimizing storage density and reliability. This promising synergy presents exciting opportunities for advancing DNA-based data storage. In future work, the applications of PERFECT PCR could be extended to include the synthesis of cost-effective, rapid, nearly error-free artificial chromosomes, mRNA vaccines, *in vitro* synthesis of biomolecules, and DNA origami.

## Author contributions

A.H., Q.S., and P.d.F. conceived the project. R.S., Q.S., and P.D. designed experiments, R.S. carried out experiments and analyzed the data. R.S contributed to writing the first draft of the manuscript. Dr. Hsing Fann, Tracy Mei, Dr. Min Feng, Siddhi Kotnis, Daniela Ruiz, Sofia Feliciano and Siddhant Gulati contributed technical support to the project. All authors revised and edited the manuscript. All authors approved the final version of the article.

## Data and materials availability

All data needed to evaluate the conclusions in the paper are present in the paper and/or the Supplementary Materials.

